# White matter integrity and nicotine dependence in smokers: evaluating vertical and horizontal pleiotropy

**DOI:** 10.1101/2020.12.09.417899

**Authors:** Zhenyao Ye, Chen Mo, Kathryn Hatch, Song Liu, Si Gao, Elliot Hong, Peter Kochunov, Shuo Chen, Tianzhou Ma

**Affiliations:** Maryland Psychiatric Research Center, Department of Psychiatry, School of Medicine, University of Maryland, Baltimore, Maryland, United States of America; Division of Biostatistics and Bioinformatics, Department of Epidemiology and Public Health, School of Medicine, University of Maryland, Baltimore, Maryland, United States of America; Department of Epidemiology and Biostatistics, School of Public Health, University of Maryland, College Park, Maryland, United States of America; Department of Computer Science and Technology, Qilu University of Technology, Jinan, Shandong, China

## Abstract

Smoking is a heritable behavior and nicotine dependency is complex mechanism supported by both positive and negative reinforcements. We hypothesized that cerebral white matter (WM) may mediate the individual dependency on nicotine integrity because its integrity is altered in smokers and shows dose-related response to nicotine administration. Two vertical and one horizontal pleiotropy pathways that combined individual genetic variations, measure of WM integrity by fractional anisotropy (FA), and nicotine dependence were evaluated in a large epidemiological sample (N=12,264 and 4,654 participants that have genetic, FA measure and nicotine dependence data available for smoking status and cigarettes per day (CPD), respectively) collected UK Biobank. We started by selecting the candidate genetic regions including genetic risk factors associated with smoking from genome-wide association study (GWAS) for causal pathway analysis. Then we identified pleiotropic loci that influence both nicotine dependence and WM integrity from these regions. We tested a horizontal pleiotropy pathway: (A) genetic risk factors associated with smoking were independently affecting both nicotine dependence and WM integrity. We also evaluated two vertical pleiotropy that assumed that individual genetic factors associated with nicotine dependence impacted B) impacted WM integrity which in turn led to higher nicotine dependence vs. C) led to nicotine dependence and resulting white matter alterations. There were 10 and 23 candidate pleiotropic variants identified for smoking status and CPD traits. All these variants exhibited vertical pleiotropy. For smoking status, the genetic effect on smoking status was mediated by FA measures over multiple brain regions. The variants were located in a gene *SARDH*, which catalyzes the oxidative demethylation of sarcosine that plays a role in reducing tolerance effect on nicotine. Conversely, CPD was a significant mediator in the vertical pleiotropy pathway to FA. The identified variants were located in gene *IREB2*, that was reported as a susceptibility gene for both neurodegeneration and smoking-induced diseases.

## Introduction

Nicotine dependence is a complex behavior and tobacco smoking is the main modifiable risk factor for cancer and aging disorder and a chief co-morbidity for mental illnesses. Population and epidemiological studies suggest strong genetic susceptibility to nicotine dependence with additive genetic factors explaining up to ~75% of the variance on nicotine-related traits. Genome wide association studies (GWAS) localized and replicated multiple genetic variants conferring susceptibility to nicotine dependence including these regulating acetylcholine receptors (nAChRs) [1–4]. The neuroanatomical mechanism of nicotine dependence remain unknown. Neuroimaging studies reported both observational [5–7] and direct associations [8] between smoking/nicotine dependence and administration and white matter (WM) integrity, assessed by the fractional anisotropy (FA) of water diffusion in diffusion tensor imaging (DTI) [9]. Chronic smoking is associated with significantly reduced FA values [10–14]. Conversely, transient elevation of FA were observed following nicotine administration in both smokers and controls [8] and higher FA values were reported in lightly smoking individuals [15] or adolescent smokers [10, 11]. Transient changes in WM integrity following nicotine administration in smokers were correlated with improvements in cognitive measures such as attention and processing [8], while lower FA values in chronic smokers were associated with lower cognitive scores [10–14]. We hypothesize that changes in cerebral WM integrity in smokers may contribute to neuroanatomic mechanisms of maintaining the nicotine addiction.

WM integrity maybe directly involved in the genetics of the positive and negative mechanisms that sustain the nicotine addiction. The positive (smoking to feel good) reinforcement of nicotine addiction is associated with cognitive and mood enhancement effects of nicotine and could be driven through, or intimately involved with the transient elevation in WM integrity following nicotine administration [8]. The negative reinforcement (smoking not to feel bad) of nicotine addiction through cognition and mood may also be caused by reduction in white matter integrity due to cardio-and-cerebrovascular risks of chronic smoking [10–14]. We propose to use mediation analysis as a tool to evaluate the involvement of cerebral WM in the mechanism of addition. On one hand, the basic genetic building blocks of addiction can act by producing a pattern of neuroanatomical changes throughout the brain, thus reinforcing addiction behavior [16]. On the other hand, the negative health consequences of chronic smoking behavior may in turn imprint the deficit patterns on brain structure and function through multiple health hazards associated with smoking.

We hypothesized three competing vertical and horizontal pleiotropy pathways to probe the relationship between genetics, WM integrity and nicotine dependence. Vertical and horizontal pleiotropy are the main causal mechanisms in complex polygenic behavior [17–19]. In horizontal pleiotropy, two traits are influenced by the genetic factors independently (Fig 1: Model 0). Vertical pleiotropy is observed when trait influenced by genetic factors in turn influences another trait by acting a mediation. In the first pathway, we hypothesize that genetic factors underwrite susceptibility to smoking severity through WM integrity (Fig 1: Model 1). In the second pathway, we alternatively hypothesize that genetic factors underwrite nicotine addiction directly, and the reduction in WM integrity occurs due to multitude of harmful effects of chronic or heavy smoking (Fig 1: Model 2). We then develop rigorous analytical approach to determine and validate the best causal model for each variant associated with both WM integrity and nicotine addiction behavior.

**Figure 1.**
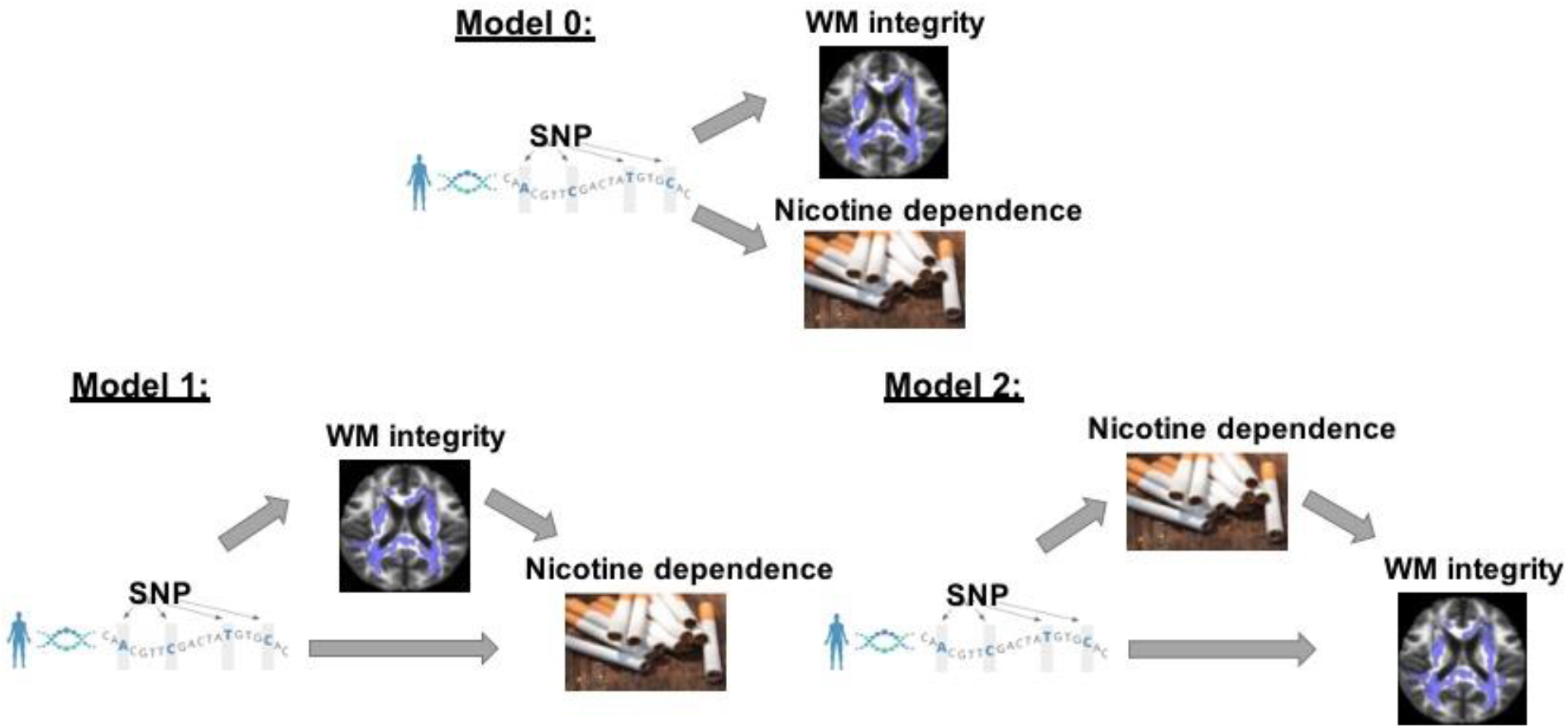
Three competing vertical and horizontal pleiotropy pathways proposed to understand the causal relationship between genetics, white matter integrity and nicotine dependence. Model 0 represents a horizontal pleiotropic relationship, while model 1 and 2 represent vertical pleiotropic relationship.

We used two smoking behavior traits smoking status and cigarettes per day (CPD) as measures of nicotine dependence to test the proposed causal pathways in a large scale epidemiological data from UK Biobank. We first performed GWAS on each trait and selected smoking behavior associated genetic regions for causal pathway analysis. We then identified 10 and 20 pleiotropic loci that influence both WM integrity and nicotine dependence for smoking status and CPD, respectively. Our data showed that WM integrity is not independent of nicotine dependence after conditioning on the genetic effect of any variants, so the relationship is not horizontal pleiotropic and we no longer proceed with model 0. Model 1 and model 2 with vertical pleiotropic relationship are essentially two alternative mediation models, thus we applied mediation analysis to select the best model. The genetic effects of the 10 variants on smoking status were mediated by FA measures in multiple brain regions. On the contrary, for the 20 variants associated with both CPD and FA, CPD acts as a mediator that mediate the genetic effects on FA. The identified variants mainly reside in two genes *IREB2* and *SARDH*, both are related to the smoking induced mechanism inside the brain and will need to be further examined for their functionality.

## Materials and Methods

### UK Biobank cohort

The data used to test our causal pathways are from the UK Biobank, a large prospective study that recruited 500,000 participants aged between 40-69 years in 2006-2010 in 22 assessment centers throughout the UK. UK Biobank data consists of phenotypic, genotypic, and imaging details about its participants collected from questionnaires, physical measures, multimodal imaging, genome-wide genotyping, and longitudinal follow-up for health-associated outcomes [20]. We restrict our genome-wide association analysis (GWAS) to include only participants with white ethnicity backgrounds (British, Irish, and any other white background) and with both genotype and smoking behavior phenotype data available. For causal pathway analysis, we further narrow down to participants who have genotype, nicotine dependence phenotype and white matter integrity phenotype data available. Number of participants included at each analytic step is summarized in the Supplement (Figure S1).

### Smoking behavior related phenotype

We chose to analyze the following two smoking behavior traits due to their relevance to nicotine dependence. Table S1 summarizes the number of participants by smoking-related phenotype codes in UK Biobank.

1. Smoking status (binary trait: current vs. never smokers). Current and never smokers were defined using phenotype codes 20116 (smoking status) in UK Biobank.
2. Cigarettes per day (CPD; average number of cigarettes smoked per day by participants who are either current or past smokers), which has been broadly used in previous studies for their relevance to nicotine addiction behavior [4–6, 21]. CPD was defined using phenotype codes 2887 (number of cigarettes previously smoked daily), 3456 (number of cigarettes currently smoked daily), and 6183 (number of cigarettes previously smoked daily (current cigar/pipe smokers)) in UK Biobank. The CPD values of participants who smoked less than one cigarette per day was recoded to 0; and the CPD values of those who smoked more than 60 cigarettes per day were recoded to 60.

### Genotype data and genome-wide association study

UK Biobank cohort was genotyped using Axiom Biobank Arrays analyzing up to over 90 million single nucleotide variants (SNVs) of 487,409 subjects. We first removed variants with minor allele frequency (MAF) below 0.01 and Hardy-Weinberg equilibrium P-value below 0.001, and removed individuals with more than 5% missing genotypes. The preprocessing step left us with 8,521,984 SNVs for 459,228 subjects. We then performed GWAS to find smoking behavior associated loci using PLINK 1.9 [22]. Most significantly associated peaks including SNPs reported in previous GWAS of smoking behavior and validated in our study were selected to carry out the causal pathway analysis. In addition, considering the strong linkage disequilibrium (LD) of neighboring SNPs, we also included all SNPs in the genomic regions +/− 250 kb around the peak boundaries.

### White matter integrity phenotype data

The UK Biobank consists of multi-modal braining imaging data covering structural, functional and diffusion imaging. In this study, we concentrate on the white matter fractional anisotropy (FA) measure derived from diffusion MRI data, a common measure of white matter integrity whose association with smoking addiction behavior has been reported in previous studies [23]. UK Biobank database provides FA measures from multiple brain regions (Table S2), including ICP, GCC, BCC, SCC, FX, CST (mean/right/left), ALIC (right/left), PLIC (right/left), RLIC (right/left), ACR (right/left), SCR (right), SCR (left), PCR (right/left), PRT (right/left), SS (right/left), EX (right/left), CGC (right/left), CHG (right/left), FXST (right/left), SLF (right/left), SFO (right/left), UN (right/left), and TAP (right/left).

### The three pleiotropic pathways

Denote by G the genotype, M the FA measures, Y the smoking behavior and Z the potential confounding covariates. Given the directed graph structures in Figure 1, we can represent the three competing models by factorizing their joint distributions:

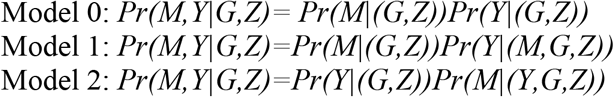

Model 0 assumes genetics to be a common cause of both FA and nicotine dependence behavior independently (“SNP->Smoking”, “SNP-> FA” and FA⊥Smoking given SNP and confounders) and represents a horizontal pleiotropic relationship. Model 1 and model 2 are two alternative mediation models and represent vertical pleiotropic relationship. In model 1 (“SNP-> FA->Smoking”), FA measures are regarded as the mediators that mediate the effect of SNPs on the nicotine dependence behavior. In contrast, model 2 (“SNP->Smoking->FA”) considers the long-term effect of chronic smoking on the brain structure and regards nicotine dependence behavior as the mediator that mediates the effect of SNPs on FA.

Our analytical procedures start by identifying the potentially pleiotropic variants of both FA and smoking traits. We then evaluate the association between FA and smoking traits given the SNP effects to distinguish horizontal pleiotropy from vertical pleiotropy. For variants with vertical pleiotropic relationship, we further conduct causal mediation analysis to choose the best mediation model that explains the relationship between SNP, FA and smoking. Below, we describe the analytical approaches in details.

#### Step 1. Identification of pleiotropic variants

Suppose we start with a set of *g*_0_ SNPs from GWAS results for n subjects, let G_ij_ denote the genotype of ith subject in jth SNP (1 ≤ *i* ≤ *n*, *j* ∈ *g*_0_), M_i_ denote the continuous FA measure, Y_i_ denote the smoking phenotype (either continuous or binary) and Z_i_ denote the covariates of ith subject. We assume an additive genetic model and let G_ij_=0, 1, or 2 represent the number of copies of minor alleles. In the first step, we look for SNPs that are associated with both FA measures and smoking traits, i.e. potentially pleiotropic variants. Note that this step is also a necessary condition to establish mediation for both model 1 and model 2, where the mediator and the outcome are simply switched in the two models [24–26]. We fit linear or logistic regression model for FA measure and smoking trait respectively on each SNP adjusting for the potential confounding covariates such as gender and age:

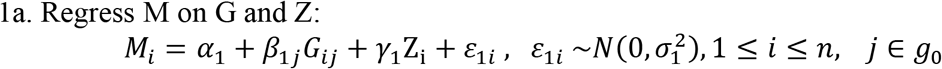

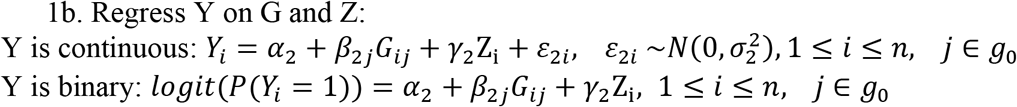

where α_1_ and α_2_ are the intercepts, γ_1_ and γ_2_ are the effects of covariates, β_1*j*_ and β_2*j*_ correspond to the genetic effect of *j*th SNP on M and Y. The cutoff for statistical significance is chosen to control for the overall false discovery rate (FDR) in identifying the potentially pleiotropic variants in the shared subset that meet both SNP-FA and SNP-smoking association criteria (see Supplement). In case of small or unbalanced sample size, looser cutoff based on p-values for each association can also be applied.

#### Step 2: Distinguish horizontal from vertical pleiotropy

Model 0 assumes horizontal pleiotropic relationship while model 1 and 2 assume vertical pleiotropic relationship. Main feature of horizontal pleiotropy is that the two traits are independent given the SNP effect. In this step, we conduct association analysis between FA and smoking traits conditioning on the SNP. If the conditional independence holds, the variants demonstrate horizontal pleiotropy; otherwise, they demonstrate vertical pleiotropy.

#### Step 3: Selection of the best mediation model for vertical pleiotropy

Model 1 and model 2 are two alternative mediation models under the vertical pleiotropy assumption. Mediation analysis investigates how a third variable affects the relation between two other variables and is a useful tool in discovering the hidden mechanism in many biological fields. We conduct exploratory mediation analysis and select the best mediation model for each pleiotropic SNP using Bayes Factor criteria. We then validate the selected model by checking the major causality assumptions and test and categorize the mediation effects.

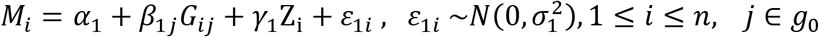

Suppose the set of SNPs that meet the criteria of vertical pleiotropy from step 1 and 2 is *g*_1_, to determine the best mediation model from the two candidates, we regress the outcome on both exposure variable and mediator for model 1 (outcome=smoking) and model 2 (outcome=FA) respectively adjusting for covariates:

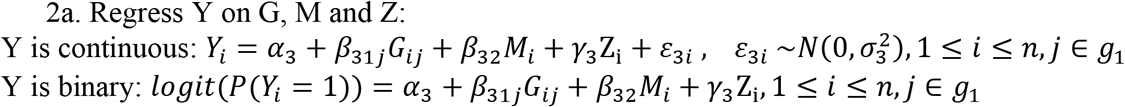

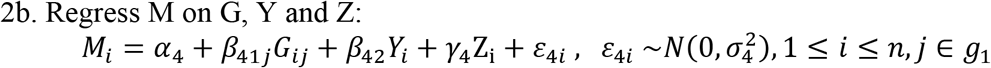

where α_3_ and α_4_ are the intercepts, γ_3_ and γ_4_ are the effects of covariates, β_31*j*_ and β_41*j*_ represent the direct effects of SNPs on outcomes Y and M in model 1 and 2, respectively, β_1*j*_β_32_ and β_2*j*_β_42_ represent the indirect effects of SNPs on outcome via the mediators M and Y in model 1 and 2, respectively.

To select the mediation model that best explains the causal relationship for each SNP j, we propose to use the penalized likelihood BIC score as a model selection criteria [27, 28]. By definition, 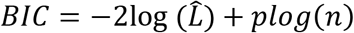, where 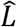 is the maximized value of the likelihood, *p* is the number of parameters and *n* the sample size. Since the two mediation models have exactly the same number of parameters, we are essentially comparing the maximum likelihoods of the two models. The maximum likelihoods of model 1 and model 2 can be derived from their joint distribution combining step 1 and step 2 (Model 1: 1a+2a; Model 2: 1b+2b), with the parameters evaluated at MLE:

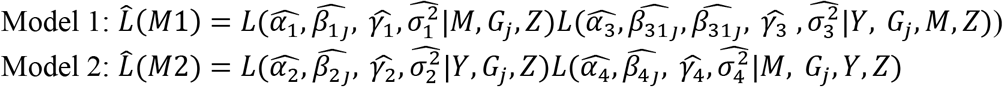

BIC can be shown to approximate the distribution of the data P(D|M) by integrating out the parameters using Laplace's method (see derivation in the Supplement), thus we can directly compute the Bayes Factor (BF), a likelihood ratio of the marginal likelihood of two competing models defined as:

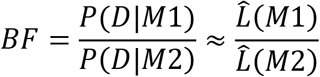

When BF>1, model 1 is preferred, otherwise model 2 is preferred. We followed from Kass and Raftery [29] to interpret BF for its strength of evidence.

The BF based model selection performs exploratory mediation analysis to determine the favored mediation model for each potentially causal variant. We will need to validate the mediation model selected by carefully checking the causal mediation assumptions (see Supplement). Once the model assumptions are checked, the causal mediation has been established. We will then test for the mediation effect in the causal path by using nonparametric bootstrap procedure [30, 31]. For model 1, β_31*j*_ represents the direct effect, β_1*j*_β_32_ represent the indirect/mediation effect and β_2*j*_ represent the total effect. For model 2, β_41*j*_ represents the direct effect, β_2*j*_β_42_ represent the indirect/mediation effect and β_1*j*_ represent the total effect. The estimated proportion of mediation effects can be computed as 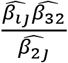 and 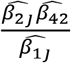 for model 1 and model 2, respectively.

Zhao et al. (2010) [32] classified mediation into three types according to significance and direction of direct effect when mediation effect is significant: when the direct effect is also significant and has the same sign, the mediation is called a complementary mediation; if they point to the opposite directions, the mediation is called a competitive mediation; lastly, if the direct effect is not significant, the mediation is indirect-only mediation. We will follow this classification to interpret the final mediation results for each variant.

All statistical analyses were conducted using R program [33]. For model checking and testing of direct and indirect effect in mediation analysis, we use the R package “mediation” [34].

## Results

### GWAS and selection of smoking associated loci

GWAS were conducted separately for smoking status (N=294,735) and CPD traits (N=142,752). Numerous important smoking behavior associated loci previously reported were reproduced in our study [4, 35–40] as highlighted in the circular Manhattan plots (Figure 2). Notably, the significant loci identified for each of the two traits have little overlap, implying the different genetic basis of the two traits of smoking behavior. The loci associated with smoking status are mainly located in regions on chromosome 9, 10 and 11 marked by the genes FAM163B, SARDH, CNNM2, NCAM1 and non-coding RNA LOC101928847, while loci associated with CPD are located in regions on chromosome 8, 15 and 19 marked by the genes CHRNB3, CHRNA3, IREB2 and RAB4B (Table 1). The results validated the gene findings of smoking behaviors in previous GWAS studies [4, 41, 42]. We will perform causal pathway analysis separately on the highly significant and reproducible loci associated with smoking status and CPD. Considering the strong linkage disequilibrium (LD) among nearby loci, we also included loci in the extended genomic regions by 250kb both up and down-stream (Supplementary Figure S3-S8). Table 1 shows the genomic regions selected including a total of 4224 SNPs for smoking status and 5828 SNPs for CPD, respectively. Next, we focus on participants who have genotype, FA measures and smoking behavior phenotype data available (N=12,264 for smoking status and N=4,654 for CPD) and perform causal pathway analysis on the potentially pleiotropic variants.

**Figure 2.**
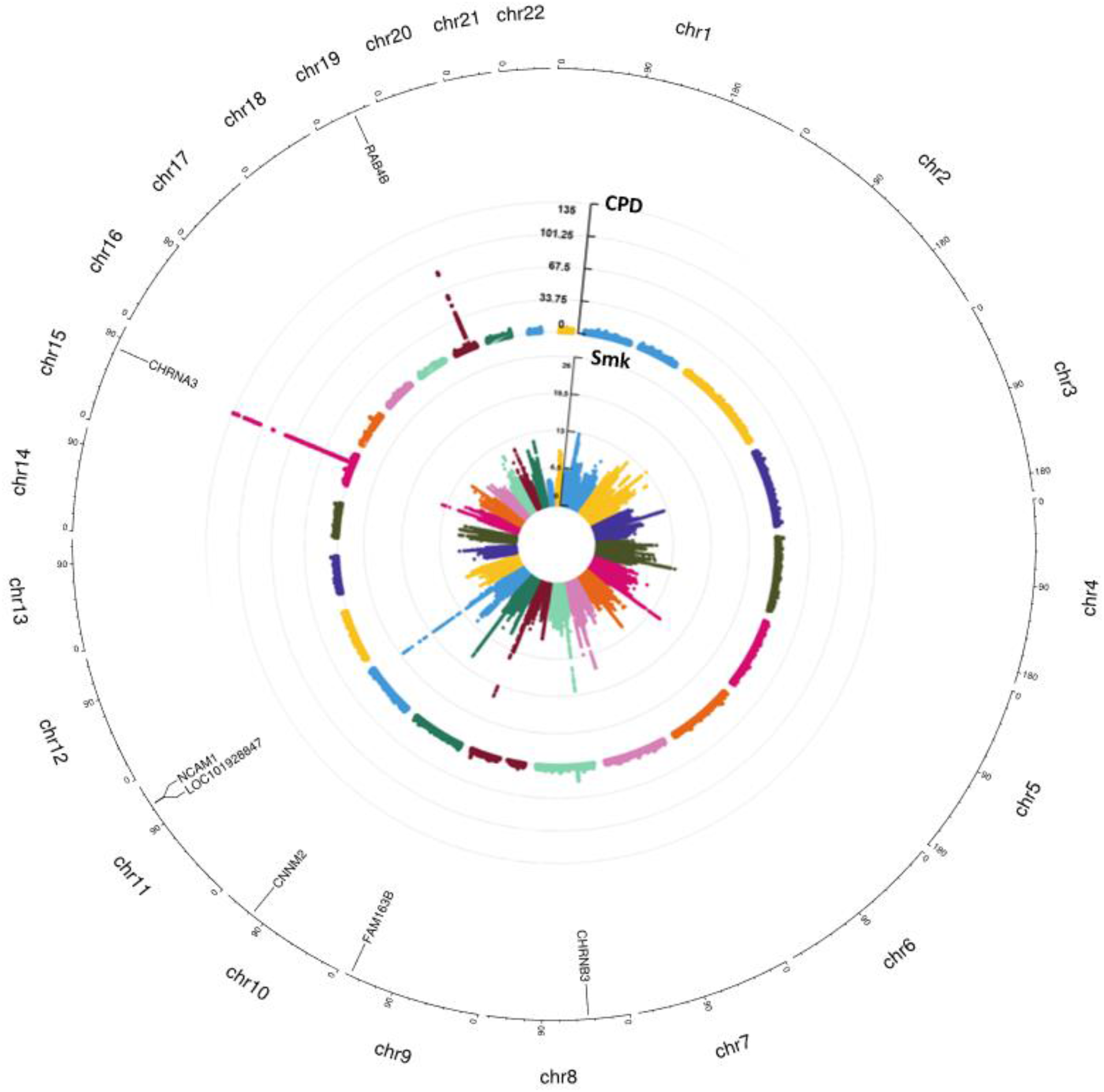
A concentric circular Manhattan plots of the GWAS results for smoking status and CPD for chromosomes 1-22. Each dot represents an SNV, X and Y axes refer to genomic locations and −log10(p-value).

**Table 1.**
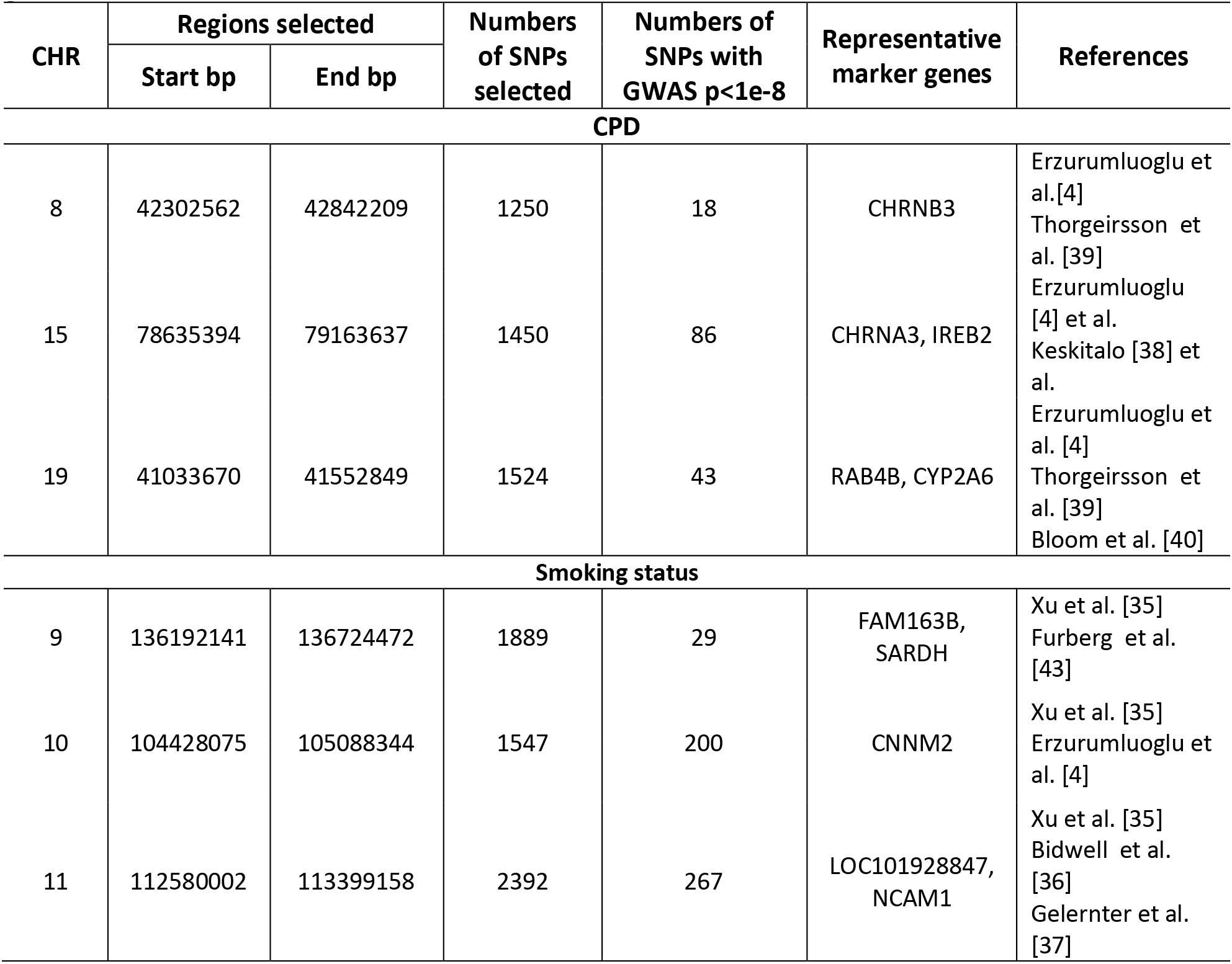
The selected genomic regions for causal pathway analysis marked by representative genes

### Causal pathway analysis for smoking status

Univariate association analysis found 28 FA measures from various brain regions that show significantly higher FA among never smokers than current smokers (p<0.05; Table S3), supporting the findings in previous literature [5, 44, 45]. We focus on these 28 FA measures for our causal pathway analysis for smoking status. There are 10 potentially pleiotropic SNPs identified that have significant effects on both smoking status and 16 FA measures (p<0.05 for both associations; Table S4). These include tracts in corpus callosum (GCC, BCC and SCC), corona radiata (ACR, SCR and PCR), sagittal stratum (SS) and posterior thalamic regions (PTR). The significant association between these FA measures and smoking status held given the genetic effects of any of the SNPs implying a vertical pleiotropic relationship (Table S5), so we no longer proceed with model 0. In comparing the two mediation models of vertical pleiotropy, model 1 where FA mediates the genetic effect on smoking status is favored (BF≫100 indicating decisive evidence [29]). Both direct and indirect effects are significant while pointing to the same directions, classified as complementary mediation effects. These variants show strong LD correlation and are located in *SARDH* gene, a gene that catalyzes the oxidative demethylation of sarcosine and has been found related to smoking behavior in large GWAS meta-analysis (Table 2; Figure 3(A)) [43].

**Table 2.** Gene annotation and minor allele information of the 10 pleiotropic SNPs associated with both smoking status and FA measures and the 23 (change to 20??) pleiotropic SNPs associated with CPD and FA measures.

| Nicotine dependence | CHR | Gene Name | SNP | Minor Allele (1000 Genome/UKBiobank) |
| --- | --- | --- | --- | --- |
| Smoking status | 9 | SARDH | rs129891 | A |
|  | 9 | SARDH | rs129895 | G |
|  | 9 | SARDH | rs2797830 | T |
|  | 9 | SARDH | rs129893 | C |
|  | 9 | SARDH | rs129899 | T |
|  | 9 | SARDH | rs129894 | G |
|  | 9 | SARDH | rs129939 | A |
|  | 9 | SARDH | rs129898 | T |
|  | 9 | SARDH | rs129892 | C |
|  | 9 | SARDH | rs129948 | G |
| Cigarette per day | 15 | IREB2 | rs1964678 | G |
|  | 15 | IREB2 | rs4887057 | G |
|  | 15 | IREB2 | rs12593229 | G |
|  | 15 | IREB2 | rs4299116 | A |
|  | 15 | IREB2 | rs1504549 | T |
|  | 15 | IREB2 | rs12910910 | T |
|  | 15 | IREB2 | rs8043227 | G |
|  | 15 | IREB2 | rs12916801 | G |
|  | 15 | IREB2 | rs8042238 | T |
|  | 15 | IREB2 | rs8042260 | G |
|  | 15 | IREB2 | rs12903295 | G |
|  | 15 | IREB2 | rs12904234 | T |
|  | 15 | IREB2 | rs36146269 | A |
|  | 15 | IREB2 | rs4887059 | T |
|  | 15 | IREB2 | rs965604 | A |
|  | 15 | IREB2 | rs13180 | T |
|  | 15 | IREB2 | rs12899351 | C |
|  | 15 | IREB2 | rs1062980 | C |
|  | 15 | HYKK, IREB2 | rs12594711 | T |
|  | 15 | HYKK, IREB2 | rs4362358 | T |
|  | 15 | IREB2 | rs11637656 | T |
|  | 15 | IREB2 | rs12592111 | A |
|  | 15 | IREB2 | rs12903285 | G |

| Nicotine dependence | Cigarette per day |  |  |  |
| --- | --- | --- | --- | --- |
|  | CHR | Gene Name | SNP | Minor Allele<br>(1000 Genome/UKBiobank) |
| Smoking status | 9 | SARDH | rs129891 | A |
|  | 9 | SARDH | rs129895 | G |
|  | 9 | SARDH | rs2797830 | T |
|  | 9 | SARDH | rs129893 | C |
|  | 9 | SARDH | rs129899 | T |
|  | 9 | SARDH | rs129894 | G |
|  | 9 | SARDH | rs129939 | A |
|  | 9 | SARDH | rs129898 | T |
|  | 9 | SARDH | rs129892 | C |
|  | 9 | SARDH | rs129948 | G |
| Cigarette per day | 15 | IREB2 | rs1964678 | G |
|  | 15 | IREB2 | rs4887057 | G |
|  | 15 | IREB2 | rs12593229 | G |
|  | 15 | IREB2 | rs4299116 | A |
|  | 15 | IREB2 | rs1504549 | T |
|  | 15 | IREB2 | rs12910910 | T |
|  | 15 | IREB2 | rs8043227 | G |
|  | 15 | IREB2 | rs12916801 | G |
|  | 15 | IREB2 | rs8042238 | T |
|  | 15 | IREB2 | rs8042260 | G |
|  | 15 | IREB2 | rs12903295 | G |
|  | 15 | IREB2 | rs12904234 | T |
|  | 15 | IREB2 | rs36146269 | A |
|  | 15 | IREB2 | rs4887059 | T |
|  | 15 | IREB2 | rs965604 | A |
|  | 15 | IREB2 | rs13180 | T |
|  | 15 | IREB2 | rs12899351 | C |
|  | 15 | IREB2 | rs1062980 | C |
|  | 15 | HYKK, IREB2 | rs12594711 | T |
|  | 15 | HYKK, IREB2 | rs4362358 | T |
|  | 15 | IREB2 | rs11637656 | T |
|  | 15 | IREB2 | rs12592111 | A |
|  | 15 | IREB2 | rs12903285 | G |

**Figure 3.**
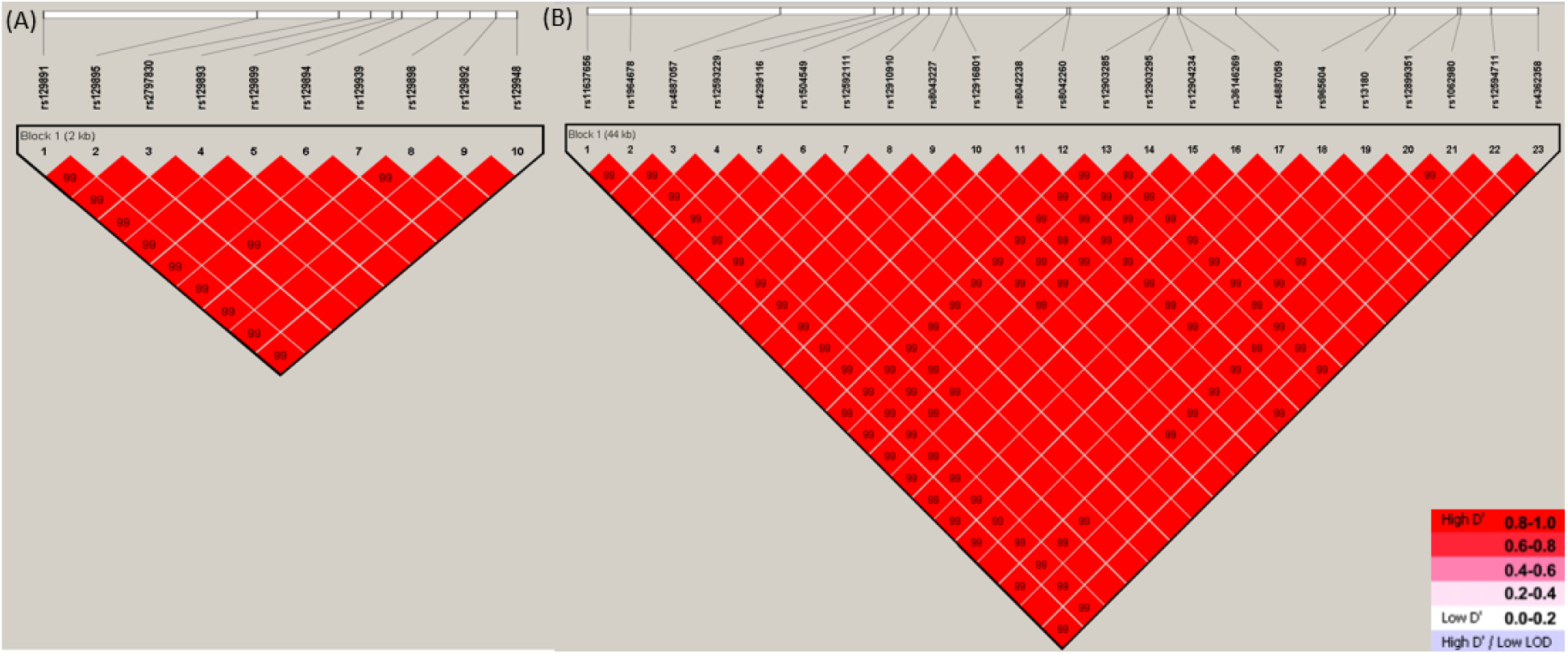
Linkage disequilibrium (LD) plot for (A) 10 SNPs associated with both smoking status and FA measures; (B) 23 (change to 20??) SNPs associated with CPD and FA measures. The values in boxes are pair-wise SNP correlations (D’), bright red boxes without numbers indicate complete LD (D’=1).

Figure 4 shows example of one variant, rs129891, to demonstrate how genetics impact smoking status via FA measure of WM. Participants with more copies of minor allele “A” are more likely to be current smokers (p=0.045). In the same time, FA measure in the three corpus callosum regions are significantly lower among individuals carrying more copies of “A” (BCC: p=4.97*10^−2^; GCC: p=1.75*10^−2^; SCC: p=3.15*10^−2^). This allele is a risk allele for both reducing white matter integrity and increasing the tendency to nicotine addiction. In the final selected mediation model, GCC, BCC and SCC act as complementary mediators with significant indirect effects (p=0.012, 0.038 and 0.018, respectively) the same direction as direct effects of SNP on smoking status (mediation proportions=3.8%, 4.5% and 6.1%, respectively), implying the consistent effect of variant on brain structure and smoking status related to neurodegeneration and smoking-induced diseases [44–47]. The causal pathways of the genetic variants involve multiple mediators of FA measures in the three corpus callosum regions highly correlated with each other (*ρ*=0.82, 0.57 and 0.51, respectively), whether they sit in parallel pathways or carry out effects in sequential order need to be further validated in future studies [19, 48, 49].

**Figure 4.**
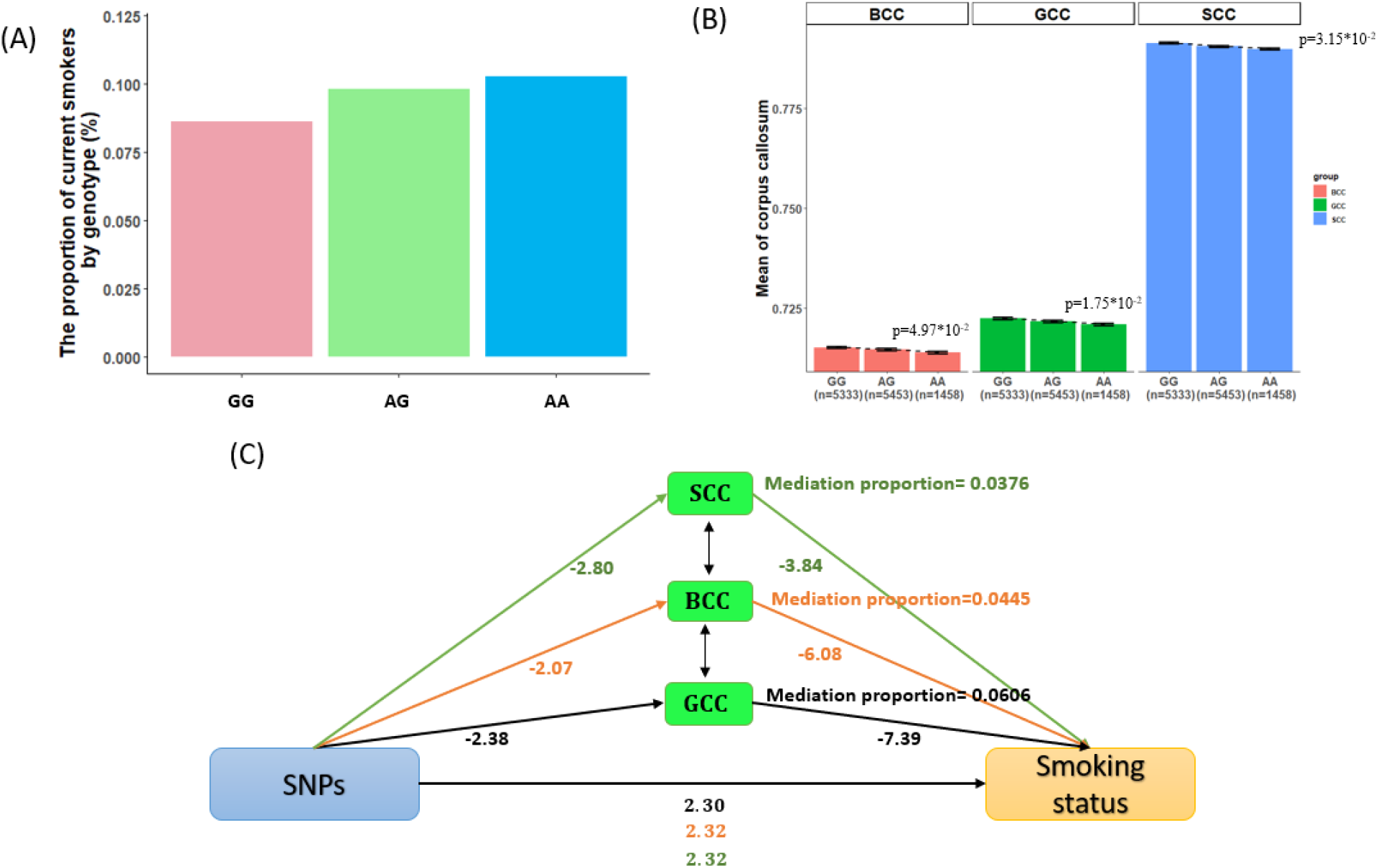
Effect of the rs129891 polymorphism on smoking status, white matter FA measures (one of the FA measures, genu of corpus callosum (GCC) as an example), and mediation model 1, respectively, in 12,264 white ethnic background participants. Panel A shows the proportion of currents smokers according to rs129891 genotype. Panel B shows mean of GCC values according to rs129891 genotype. Sample sizes for each genotype group are shown in parentheses under the horizontal axis. Panel C shows the mediation effects 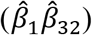 of rs129891 is 17.59,12.59 and 10.75 in mediation model separately with GCC, BCC and SCC as mediators and smoking status as an outcome.

### Causal pathway analysis for CPD

Univariate association found 21 FA measures that show significant negative association with CPD (p<0.05; Table S3). We focus on these 21 FA measures for our causal pathway analysis for smoking status. A total of 20 pleiotropic SNPs were identified with significant effects on both CPD and 7 FA measure (overall FDR<0.15; Table S4). These include tracts in corpus callosum (GCC and BCC), corona radiata (SCR and PCR) and internal capsule (RLIC). The significant association between these FA measures and smoking status held given the genetic effects of any of the SNPs implying a vertical pleiotropic relationship (Table S5), so we no longer proceed with model 0. In comparing the two mediation models for vertical pleiotropy, interestingly, all the 20 variants favor model 2, where CPD mediates the genetic effect on FA (BF≫100 indicating decisive evidence [29]). Both direct and indirect effects are significant but pointing to the opposite directions, classified as competitive mediation effects. All these variants show strong LD with each other and are located within the iron-responsive element binding protein 2 (*IREB2*) gene (Table 2; Figure 3(B)). *IREB2* binds to iron-responsive elements to regulate iron mechanism in human and has been reported to be a susceptibility gene for both.

We use a SNP example, rs13180 (minor allele: T), to illustrate how genetics influence FA measures indirectly via CPD in mediation model 2 (Figure 5). The mean values of both CPD (p= 8.77*10^−5^) and FA measure (p=5.99*10^−5^) are significantly higher among individuals with more copies of minor allele “T”. This allele plays a protective role with inherently higher white matter integrity while in the same time being a risk allele increasing the tendency to nicotine addiction. CPD is acting as a competitive mediator with indirect effect (p=0.012) of an opposite direction from the direct effect of SNP on FA, implying the adverse of effect CPD on brain structure related to neurodegeneration and smoking-induced diseases [46, 47].

**Figure 5.**
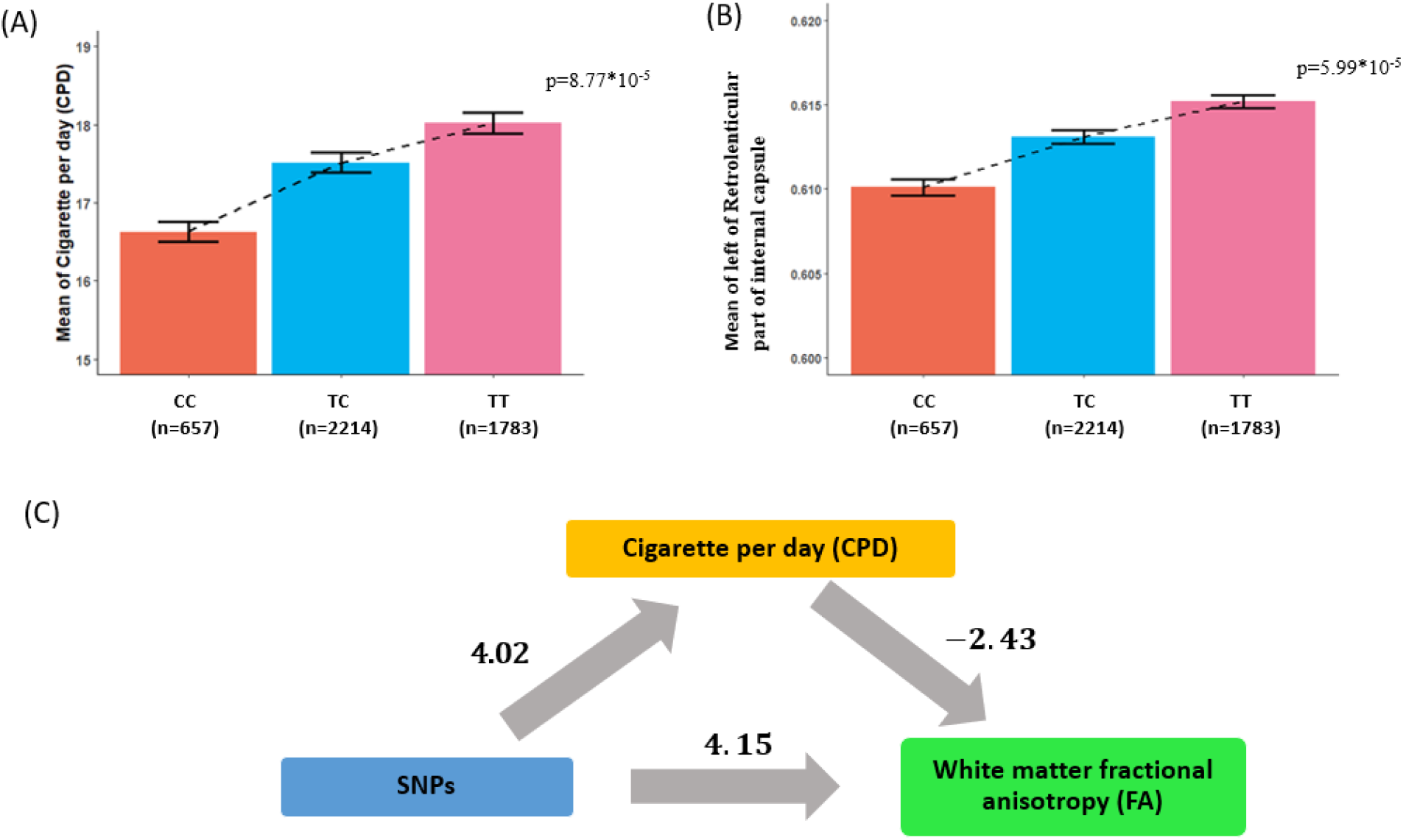
Effect of the rs13180 polymorphism on CPD, white matter FA measures (one of the FA measures, left of Retrolenticular part of internal capsule (RLIC.L) as an example), and mediation model 2, respectively, in 4654 white ethnic background participants. Panel A shows mean of CPD values according to rs13180 genotype. Panel B shows mean of RLIC.L values according to rs13180 genotype. Sample sizes for each genotype group are shown in parentheses under the horizontal axis. Panel C shows the mediation effects 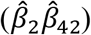 of rs13180 is −9.75 in mediation model with CPD as a mediator and RLIC.L as an outcome.

## Discussion

Enduring smoking has been found to be highly heritable, but the mechanism of nicotine addiction remain unknown. In light of the associations between nicotine dependence and white matter integrity previously reported, in this study, we performed imaging-genetic analysis for smoking status and CPD and tested three competing causal models to explain the complex causal relationship among genetics, WM integrity and nicotine dependence. Our GWAS analysis validated many reliable and reproducible loci previously reported associated with smoking related traits. We then performed causal pathway analysis on these loci and found 10 potentially pleiotropic SNPs associated with 16 FA measures in varying brain regions and smoking status, and 20 SNPs associated with 7 FA measure in RLIC (L) and CPD. The 10 SNPs for smoking status demonstrate vertical pleiotropy and favor mediation model 1 where FA mediates the genetic effect on smoking status. They are located in *SARDH*, a gene that catalyzes the oxidative demethylation of sarcosine to reduce tolerance effect on nicotine. On the other hand, the 20 vertical pleiotropic SNPs for CPD favor mediation model 2 where CPD mediates the genetic effect of these SNPs on FA measure in GCC, BCC, SCR, PCR and RLIC regions and located in *IREB2*, which regulates iron mechanism in the cell and is a susceptibility gene for both neurodegeneration and smoking-induced diseases. The basic genetic components of addiction might have produced a pattern change in WM among smokers, reinforcing the addiction behavior. Chronic severe smoking (reflected in e.g. CPD) will have negative impact to overall health which in turn reduce the WM integrity. These results enlighten the nicotine addiction study and improve our understanding of the mechanism of nicotine dependence. Further validation in a large independent cohort is needed to confirm these causal analysis results and for a more comprehensive understanding of addiction behavior.

In this study, we have used an imaging-genetics approach to study nicotine addition mechanism. Traditional imaging genetics treat neuroimaging traits in the brain as intermediate phenotype sitting along a mechanistic pathway through which genetic variation affects clinical or behavioral phenotypes. Such unidirectional model has gradually become a convention in imaging genetics mediation studies, though the justification is largely based on theory without model checking or evidence support from the data [50]. As the new technology develops and data and knowledge continue to grow, new models are in need to explain the complex interplay among genetics, brain and behavior. In this article, we have first proposed competing horizontal pleiotropic and two alternative vertical pleiotropic models assuming that the long-term behavior phenotype can mediate the genetic effect on imaging and developed a rigorous mediation analytical approach to guide users select the best model. For vertical pleiotropic model selection, we essentially perform exploratory mediation analysis to select the favored mediation model with larger BF and then confirm the selected model by checking the necessary assumptions.

Classical mediation pathway analysis has focused on analyzing univariate exposure and univariate mediator. Entering the big data era as we witness more data generated by high-throughput technology, models with multiple high-dimensional exposures and mediators have drawn more attention in the field in recent years [51–53]. Our analytical approach shows an example of performing mediation analysis when there are thousands of candidate exposure variables available in imaging genetics study. A thoughtful scheme is needed to select the relevant causal variants in the pathway while in the same time controlling for the false discovery rate in multiple comparisons. The FA measures we used in this study are summary measures over brain regions of interest using the ENIGMA work flow. For the full resolution analysis of neuroimaging trait and genotype in the causal pathway, we will consider voxel-wise neuroimaging features in future studies. More sophisticated procedures are needed for variable and model selection in such mediation analysis with potentially high-dimensional exposures, mediators and outcomes.

